# Multimodal Imaging-Based Targeting Approach for Network-Level Brain Stimulation

**DOI:** 10.64898/2026.01.09.698588

**Authors:** Alireza Shahbabaie, Mohamed Abdelmotaleb, Harun Kocataş, Filip Niemann, Daria Antonenko, Agnes Flöel, Marcus Meinzer

## Abstract

**Background:** Neural network effects of transcranial direct current stimulation (tDCS) are poorly understood. Here, we introduce an empirically informed, multimodal functional magnetic resonance imaging (fMRI) based framework suited for guiding stimulation target selection and hypothesis-based data analysis in focal tDCS-fMRI studies.

**Methods:** We illustrate our approach using data of 37 healthy individuals (19 females; mean ± SD age = 25.8 ± 5.9) recruited from a multicenter tDCS-fMRI study. Participants completed resting-state (RS- and task-fMRI (object-location memory, OLM, or associative picture-pseudoword learning, APPL, experiments) with placebo tDCS. Seed-based RS data analysis identified the functional networks originating from project-specific target regions for focal tDCS (right occipito-temporal cortex, rOTC; left ventral IFG, lvIFG) and also established their test-retest reliability using intraclass correlation coefficients (ICC). Dice coefficients analyzed the overlap between the seeded RS- and task-evoked networks. This aimed to identify task-active regions potentially affected by downstream neural network effects from the target regions.

**Results:** Seed-based analyses identified two highly reliable ventral visual-limbic (rOTC) and language-related networks (lvIFG), with >72-77% of voxels showing good-to-excellent TRR (ICC ≥ 0.75). Only a subset of voxels identified by the RS analyses overlapped with activity elicited by the experimental paradigms (ranging from 7.5-55%), with larger correspondence for the OLM task (Dice OLM: 0.249-0.349; APPL 0.065-0.106). Therefore, the degree of potential tDCS network effects varied substantially depending on the target region, the extent of its functional network and task-specific activity patterns. Degree of correspondence was further mediated by the selected contrasts-of-interest in the task-based analyses, with more conservative control conditions resulting in reduced overlap.

**Conclusion:** We established a principled, multimodal fMRI framework bridging a critical gap in neuromodulation research. By integrating reliable intrinsic connectivity maps with task-evoked activity patterns, we provide a method to prospectively identify network-level targets for focal brain stimulation and to generate hypotheses for data analyses in tDCS-fMRI studies. This approach paves the way for investigating modulation of specific functional networks, shifting the rationale from stimulating an isolated brain region to strategically targeting key nodes within a predefined functional pathway.

## 1. Introduction

Transcranial direct current stimulation (tDCS) applies weak electrical currents via scalp-attached electrodes to modulate excitability and plasticity in the human brain (Stagg 2018 J ECT) and beneficial effects of tDCS on motor and cognitive functions have been demonstrated in healthy and clinical populations (Kang et al., 2024; Lv et al., 2024; Narmashiri & Akbari, 2025). However, while the basic neurophysiological mechanisms of tDCS are well understood (Cirillo et al., 2017), its effects on the complex neural networks underlying human brain functions requires further investigation. Although carefully designed behavioral studies can provide evidence for causal brain-behavior relationships (e.g., Gbadeyan, McMahon, et al., 2016; Martin et al., 2020), they are limited to specific target regions. In addition, computational models can be used to simulate the distribution and intensity of the induced current, to investigate which brain regions are potentially affected by tDCS (Hunold et al., 2023). Therefore, combining tDCS with functional neuroimaging is essential to reveal its effects on large-scale functional brain networks and their relationship to behavioral modulation.

Functional magnetic resonance imaging (fMRI) offers high spatial and sufficient temporal resolution to study tDCS effects at the whole brain level (Esmaeilpour et al., 2020). By administering tDCS concurrently with task-based or resting-state (RS) fMRI, it is possible to investigate immediate stimulation effects on brain function. Furthermore, structural data acquired in the same session can be used for individualized current modeling and to verify accurate electrode placement (Meinzer et al., 2024; Niemann,, et al., 2024). Using this approach, several studies have demonstrated that tDCS can modulate local brain activity at the stimulation site, activity in remote areas and also large-scale functional network organization (for a review, see Ghobadi-Azbari et al., 2021). However, the vast majority of other concurrent tDCS-fMRI studies have employed conventional montages using relatively large electrodes (25-100 mm^2^), resulting in widespread current flow between electrodes, potentially affecting multiple brain regions and networks simultaneously (e.g., Saturnino et al., 2019). Consequently, the origin of the observed changes in brain function (e.g., neural network modulation vs. current spread) remained difficult to ascertain.

More recently developed focal tDCS protocols use smaller electrodes and concentric arrangement of cathodes around a center anode, to allow more precise current delivery compared to conventional montages (e.g., Ostrowski et al., 2022; Niemann, et al., 2024). This improved focality raises the possibility for network-specific neural modulation, but focal tDCS-fMRI studies are scarce (e.g., Gbadeyan, Steinhauser, et al., 2016; Lefebvre et al., 2019; Lim et al., 2024; Müller et al., 2023) and there is currently no established framework for testing hypotheses regarding network-specific neural modulation (Esmaeilpour et al., 2020; Opitz et al., 2018). To address this gap in knowledge, we developed and applied a novel analytical framework using data from the preparatory stage of a large-scale, multicenter tDCS-fMRI study (Research Unit 5429, RU; www.memoslap.de/en/home/). Specifically, we used task- and RS fMRI data from two RU projects that assessed neural correlates of object-location memory (OLM) and associative picture-pseudoword learning (APPL) tasks that were acquired with identical acquisition parameters at the same scanner. In a first step, we employed seed-based RS data analysis to identify the functional networks associated with potential target regions for tDCS in the respective projects (i.e., right occipito-temporal cortex, rOTC, in the OLM task; left ventral IFG, lvIFG, in the APPL) and also established their test-retest reliability using intraclass correlation coefficients (ICC). Second, we identified task-based activity patterns elicited by the respective learning tasks. Third, we assessed the overlap between the RS functional networks (from step one) and the task-active regions (from step two), to determine which task-based effects would represent true network-level effects of focal tDCS. Hence, we aimed to develop an empirically informed, multimodal network-based framework, suited for planning of future focal tDCS-fMRI studies and to guide hypothesis-based data analysis.

## 2. Materials and Methods

This methodological approach was applied to datasets from the MeMoSLAP research consortium. All data were acquired on the same scanner using identical acquisition parameters. The experimental setup (e.g., tasks, electrode configurations and neuronavigated placement, Niemann et al., 2024), was identical to that of subsequently planned active focal tDCS arms within the larger project, and included sham tDCS in all sessions (see section 2.1.4 for details). This approach ensured that the present data can serve as a direct comparator for future studies within the consortium. The study was pre-registered (Open Science Framework; https://osf.io/t37u2).

### 2.1. Experimental Design

#### 2.1.1. Participants

A total of 37 healthy young to middle-aged participants (19 female; mean ± SD age = 25.8 ± 5.9, range: 19-42) were included in the study. Participants were subsequently assigned to complete either the OLM task (n=18; 8 females; mean ± SD age = 25.5 ± 5.5) or the APPL task (n=19; 11 females; mean ± SD age = 26.1 ± 6.3). Interested participants were screened tDCS-fMRI eligibility via telephone and completed a comprehensive baseline assessment to ensure normal cognitive functions, followed by four combined tDCS-fMRI sessions (two RS and two task-based fMRI sessions). All participants were right-handed based on the Edinburgh Handedness Inventory (Oldfield, 1971) native German speakers, and did not report any past or current neurological or psychiatric diseases. All experimental procedures were approved by the medical ethics committee of the University Medicine Greifswald. Prior to their study inclusion, all participants provided informed consent and received monetary compensation upon study completion.

#### 2.1.2. Procedure

The four fMRI sessions were conducted at least one week apart. RS data acquisition comprised two consecutive 10-minute runs per session. Participants were instructed to keep their eyes open while fixating on a central cross-hair. Afterwards, participants were assigned to the task-based fMRI groups and completed either the OLM (n=18) or APPL (n=19) tasks.

#### 2.1.3. Cognitive task-based imaging

Presentation of the two task-related paradigm (OLM, APPL) used Presentation® software (Version 20.1, Neurobehavioral Systems, Inc., Berkeley, CA, USA) and an MRI-compatible back-projection system. Behavioral responses were recorded using MRI-compatible response grips (NordicNeuroLab, Norway; www.nordicneurolab.com). In order to minimize potential learning effects, two parallel versions of each task were administered in a counterbalanced order across sessions. Control conditions followed the same structure, but did not involve learning (see below for details).

##### 2.1.3.1. Object-Location Memory (OLM) Task

The OLM task, adapted from prior studies (Antonenko et al., 2018; De Sousa et al., 2020; Flöel et al., 2012; Prehn et al., 2017) and required participants to learn associations between objects (houses) and their spatial locations on a 2-dimensional map. The task employed a block design consisting of four learning stages, integrating both instruction, and feedback-based learning mechanisms. The control condition retained the same structure but was devoid of associative learning, requiring only left/right decisions about houses on the map. Further details on the intra-scanner task design are provided in Abdelmotaleb et al. (2025).

##### 2.1.3.2. Associative Picture-Pseudoword Learning (APPL) task

The APPL task was adapted from Sliwinska et al. (2017) and involved learning associations between pictures of common objects and pseudowords using an explicit instruction- and reinforcement-based paradigm. The experiment employed a block design, encompassing six APPL blocks and two control task blocks, with 40 trials per block. While the overall structure of the APPL and control tasks was identical, only the APPL condition involved active associative learning. Further details on the intra-scanner task design are provided in Kocataş et al. (2025).

#### 2.1.4. Focal tDCS

Focal sham tDCS was administered with a 3×1 tDCS montage using an MRI-compatible stimulator (DC-STIMULATOR MC, NeuroConn GmbH). The sham tDCS protocol involved a 10-second ramp-up to 2 mA, 20 seconds of active stimulation, and a 10-second ramp-down. Sham tDCS was started immediately prior to commencing the functional scans. Montages were individually optimized for each participant (see Niemann,, et al., 2024). The overall procedure ensured precise targeting and participant blinding, and provides a direct comparator for future active tDCS arms within the MeMoSLAP consortium.

#### 2.1.5. MRI Acquisition

All data were acquired on a 3T Siemens MAGNETOM Vida scanner with a 64-channel head-neck coil (Siemens Healthineers, Germany). Functional MRI used a multiband EPI sequence (CMRR, University of Minnesota, www.cmrr.umn.edu/multiband) with the simultaneous multislice (SMS) acceleration for high temporal resolution and whole-brain coverage (Setsompop et al., 2012). Key acquisition parameters included: 2×2×2 mm^3^ voxels, TR=1000 ms, TE=30 ms, flip angle=60°, FOV=220 mm, multiband factor=6, GRAPPA acceleration=2, and anterior-posterior phase encoding Parameters: 2×2×2 mm^3^ voxels, TR=1000 ms, TE=30 ms, flip angle=60°, FOV=220 mm, multiband factor=6, GRAPPA acceleration=2, anterior-posterior phase encoding. Resting-state sessions comprised two 10-minute runs (600 volumes each); task sessions lasted ∼30 minutes (1800 volumes). Field maps were acquired for distortion correction.

High-resolution T1-weighted MPRAGE (0.9 mm isotropic; TR=2700 ms, TE=3.7 ms, TI=1090 ms) and T2-weighted images (0.9 mm isotropic; TR=2500 ms, TE=349 ms) were acquired for structural reference and normalization of functional images.

### 2.2. Statistical Analysis

#### 2.2.1. Preprocessing

RS-fMRI data preprocessing was conducted using fMRIPrep 24.1.1 (Esteban et al., 2019), implementing a standardized pipeline for reproducible analysis. Fieldmap estimation utilized FSL’s topup based on available EPI references to correct B0 inhomogeneities. For anatomical data, T1-weighted images underwent N4 bias field correction, skull-stripping using ANTs (OASIS30ANTs template), and tissue segmentation via FSL’s FAST. Surface reconstruction was performed with FreeSurfer 7.3.2, incorporating T2-weighted data for improved pial surface estimation. Spatial normalization transformed data to both MNI152NLin6Asym and MNI152NLin6Asym:res-02 standard spaces using ANTs’ nonlinear registration.

Functional preprocessing included motion correction (FSL’s mcflirt), fieldmap-based distortion correction, and boundary-based coregistration to anatomical space (FreeSurfer’s bbregister). Confound regressors comprised framewise displacement (FD > 0.5mm threshold), the temporal Derivative of root mean square VARiance over voxelS (DVARS), anatomical and temporal CompCor components (50% variance explained), and global signals. All transformations were combined in a single interpolation using cubic B-spline sampling through nitransforms. We visually inspected the automatically generated quality control metrics at each processing stage to verify data quality.

#### 2.2.2. Confound Time Series Removal

We implemented a denoising pipeline following an optimized strategy recently suggested by Wang et al. (2023) in their systematic comparison of resting-state fMRI denoising strategies. The confound time series generated during fMRIPrep preprocessing were used to extract nuisance regressors.

Our denoising model incorporated 27 regressors, comprising a comprehensive set of 24 motion parameters (Friston et al., 1994; Satterthwaite et al., 2013) to capture both linear and nonlinear motion artifacts while maintaining orthogonality between regressors, global signal (GS), and mean signals from white matter (WM) and cerebrospinal fluid (CSF).

We automated this workflow through a custom MATLAB script (R2025a) utilizing SPM12 (version 7771) and the CONN toolbox (v22.2407, RRID:SCR_009550) with its batch processing functionality (available at [https://github.com/ShahAliR/memoslap-denoising-pipeline]) to enhance quality assurance and enable seed-based analyses within the same computational framework.

#### 2.2.3. Seed-Based Functional Connectivity

Seed regions for the RS analysis were determined by the envisaged target regions for focal tDCS, based on their established relevance for the respective learning tasks (rOTC for OLM: Gillis et al., 2016; Rolls et al., 2024; lvIFG for APPL: Perceval et al., 2020; Meinzer et al., 2012; Riemann et al., 2024) studies. Two spherical seed regions (6 mm radius) were created and centered on the MNI coordinates 47, -64, -11 (rOTC) and -47, 20, 2 (lvIFG). Binary masks for these seeds were created using Nilearn (Abraham et al., 2014) and imported into the CONN toolbox for seed-based connectivity analysis.

First-level seed-to-voxel connectivity maps were calculated by correlating the mean BOLD time series of each seed region with every other voxel in the brain, separately for each session and subject. These correlation coefficients were Fisher z-transformed to improve normality for group-level analysis. At the group level, one-sample t-tests were conducted across all 37 participants, including both sessions for each subject. This analysis tested whether the average Fisher z-transformed connectivity, aggregated over both sessions and all subjects, differed significantly from zero. Statistical significance was determined by applying a cluster-level family-wise error (FWE) correction at p < 0.05, using an initial voxel-wise threshold of p < 0.001. For each significant cluster, the peak t-value, MNI coordinates, and cluster size (k) are reported.

#### 2.2.4. Reliability Assessment

Test-retest reliability of lvIFG and rOTC functional connectivity maps was assessed using voxel-wise intraclass correlation coefficients (ICC) via Pingouin (Vallat, 2018). We computed ICC(3,k) maps using a two-way mixed-effects model for absolute agreement across sessions.

Significant clusters from seed-based connectivity analyses (FWE-corrected, p < 0.05) served as binary masks to extract ICC values within each functional network. Voxels were classified into reliability tiers (Koo & Li, 2016): poor (ICC < 0.50), moderate (0.50-0.75), good (0.75-0.90), and excellent (ICC ≥ 0.90).

#### 2.2.5. Task-based fMRI Analysis

Task-based fMRI data for the OLM and APPL paradigms were analyzed separately using SPM12 (v7771) in MATLAB R2022a. Structural T1-weighted images were skull-stripped, bias-corrected, and normalized to MNI space. Functional BOLD images were preprocessed with realignment, unwarping, slice-time correction (TR = 1 s, 72 slices), co-registration to the structural T1, normalization using the resulting deformation fields, and spatial smoothing with a 6-mm FWHM Gaussian kernel. First-level general linear models (GLMs) were constructed for each participant, incorporating data from both fMRI sessions into a single design matrix. The models included regressors for trial onsets and durations of learning and control conditions across four stages, convolved with the canonical hemodynamic response function. Six motion parameters and framewise displacement were included as nuisance regressors. Subject-specific contrast maps for the primary effect (Learning > Implicit baseline, and Learning > Control) were generated and subsequently entered into second-level one-sample t-tests. Group-level statistical maps were thresholded at p < 0.001 (voxel-level) and cluster-corrected for family-wise error (FWE) at p < 0.05. The resulting activation maps were overlaid on an MNI template, anatomically labeled using the Harvard–Oxford atlas, and visualized with MRIcroGL. More details of the task-based fMRI analyses for the OLM and APPL paradigms are provided in Abdelmotaleb et al. (2025) and Kocataş et al. (2025), respectively.

#### 2.2.6. Spatial Overlap Analysis

An identical analytical pipeline was applied to the both functional networks, wherein resting-state connectivity maps from the rOTC seed were compared to OLM task activation, and those from the lvIFG seed were compared to APPL task activation. We further considered effects of specific analytical choices for task-based activity analyses by investigating both the complex (learning > control task) and simple (learning > implicit baseline) contrasts. Spatial correspondence was quantified using the Dice similarity coefficient and voxel-wise Pearson correlation of t-values within a gray-matter mask. Analyses were conducted in Python (v3.x) using NiBabel and Nilearn. Code availability: https://github.com/ShahAliR/NetworkOverlap-Task-RS-fMRI.

## 3. Results

### 3.1. Resting-State Functional Connectivity

Whole-brain functional connectivity analyses revealed distinct large-scale networks for each seed. The rOTC seed engaged a ventral visual-limbic network, associated with integration of object recognition with contextual memory processing (Gillis et al., 2016; Kravitz et al., 2013; Rolls et al., 2024). The lvIFG seed mapped onto a left-lateralized language-semantic control network supporting lexical, phonological and conceptual integration (Fedorenko et al., 2012; Tagarelli et al., 2019). All reported clusters survived rigorous cluster-level FWE correction (p<0.05).

#### 3.1.1. Seed-Based Functional Connectivity of the rOTC

Using the rOTC as the seed region, whole-brain connectivity analysis revealed a distributed network comprising occipital, temporal, parietal, frontal, subcortical, and cerebellar regions (**Supplementary Table 1** and **Figure 1**). The most robust positive connectivity was observed with the right occipital fusiform gyrus (lateral visual stream; peak T = 17.70, cluster size = 8,973 voxels, p<0.000001 FWE-corrected) and the left occipital pole (medial early visual cortex; peak T = 11.19, cluster size = 6,682 voxels, p<0.000001). Connectivity extended to higher-order association areas, including the left supramarginal gyrus (posterior division), bilateral superior temporal gyrus, and temporal fusiform cortex. Subcortical associations included the bilateral amygdala (laterobasal and centromedial subnuclei) and parahippocampal gyrus (anterior and posterior collateral sulcus). Additional significant clusters included the cerebellar Crus I, lateral occipital cortex (superior division), and bilateral precentral and postcentral gyri.

**Figure 1.**
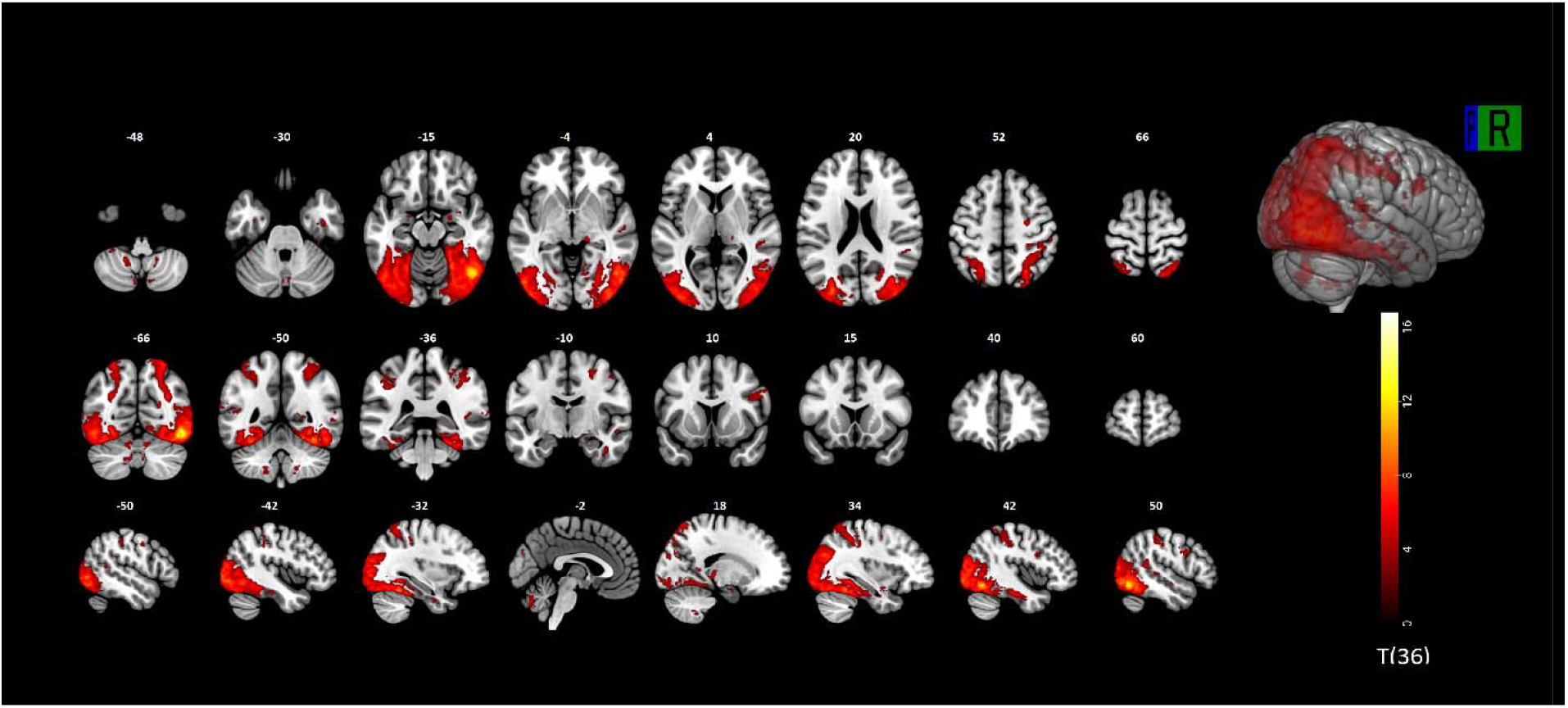
Functional connectivity of the right occipito-temporal cortex (rOTC). Whole-brain positive functional connectivity map for the rOTC seed, overlaid on the MNI152 high-resolution T1-weighted template. Connectivity strength is represented by a graded T-statistic scale from dark red (lower T-values) to bright yellow (highest T-values). Displayed clusters met voxel-wise significance at p<0.001 and survived cluster-level family-wise error (FWE) correction at p<0.05. Multiple axial, sagittal, and coronal slices are presented to capture the complete spatial distribution of the network

#### 3.1.2. Seed-Based Functional Connectivity of the lvIFG

Seeding the lvIFG, revealed a network predominantly involving temporal, parietal, and cingulo-opercular regions, as well as subcortical and cerebellar structures (**Supplementary Table 2** and **Figure 2**). The largest cluster encompassed the left superior temporal gyrus (anterior division; peak T = 20.32, cluster size = 13,145 voxels, p<0.000001) extending into adjacent auditory and language-related cortices. Significant bilateral temporal lobe connectivity included the right temporal pole (peak T = 9.26) and right middle and inferior temporal gyri. Additional strong associations were observed in the left angular gyrus (PGp), bilateral supramarginal gyrus, occipital fusiform gyrus, and precuneus. Subcortical connectivity was identified in the left thalamus (pulvinar), right thalamus (mediodorsal), bilateral caudate, and left hippocampus. Midline connectivity encompassed the anterior and mid-posterior cingulate cortices, as well as the brainstem (midbrain).

**Figure 2.**
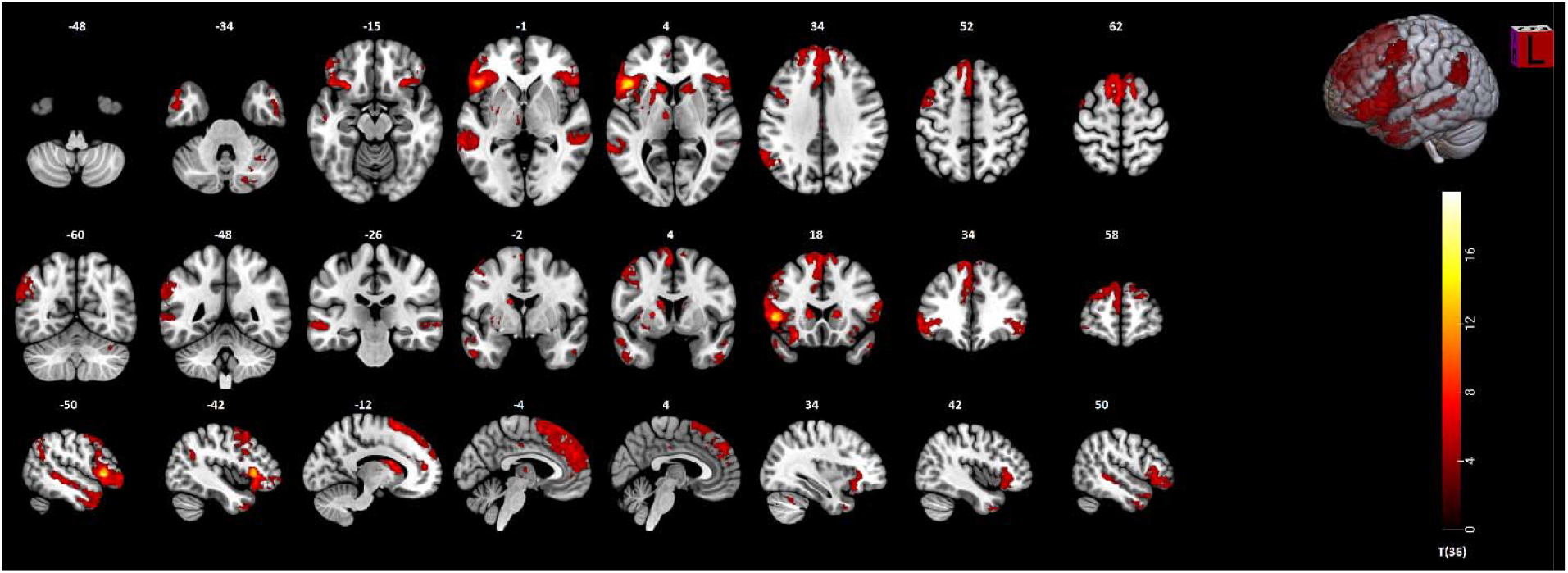
Functional connectivity of the left ventral inferior frontal gyrus (lvIFG). Positive functional connectivity network of the lvIFG seed, displayed using a red-to-yellow T-value spectrum on the MNI152 high-resolution T1-weighted template. Dark red regions indicate weaker and bright yellow indicate stronger connectivity. All clusters survived family-wise error (FWE) correction at p<0.05 (cluster-level), thresholded at voxel-wise p<0.001. Axial, coronal, and sagittal slices are shown to illustrate the full spatial extent of the network.

### 3.2. Reliability of Seed-Based Functional Networks

#### 3.2.1. Reliability of the rOTC Functional Network

The rOTC network demonstrated high test-retest reliability (**Figure 3**). The distribution of intraclass correlation coefficient (ICC) values across the 16,067 voxels within the network had a mean of 0.70 (SD = 0.14) and a median of 0.74. The interquartile range (IQR) spanned from 0.64 (25th percentile) to 0.76 (75th percentile).

**Figure 3.**
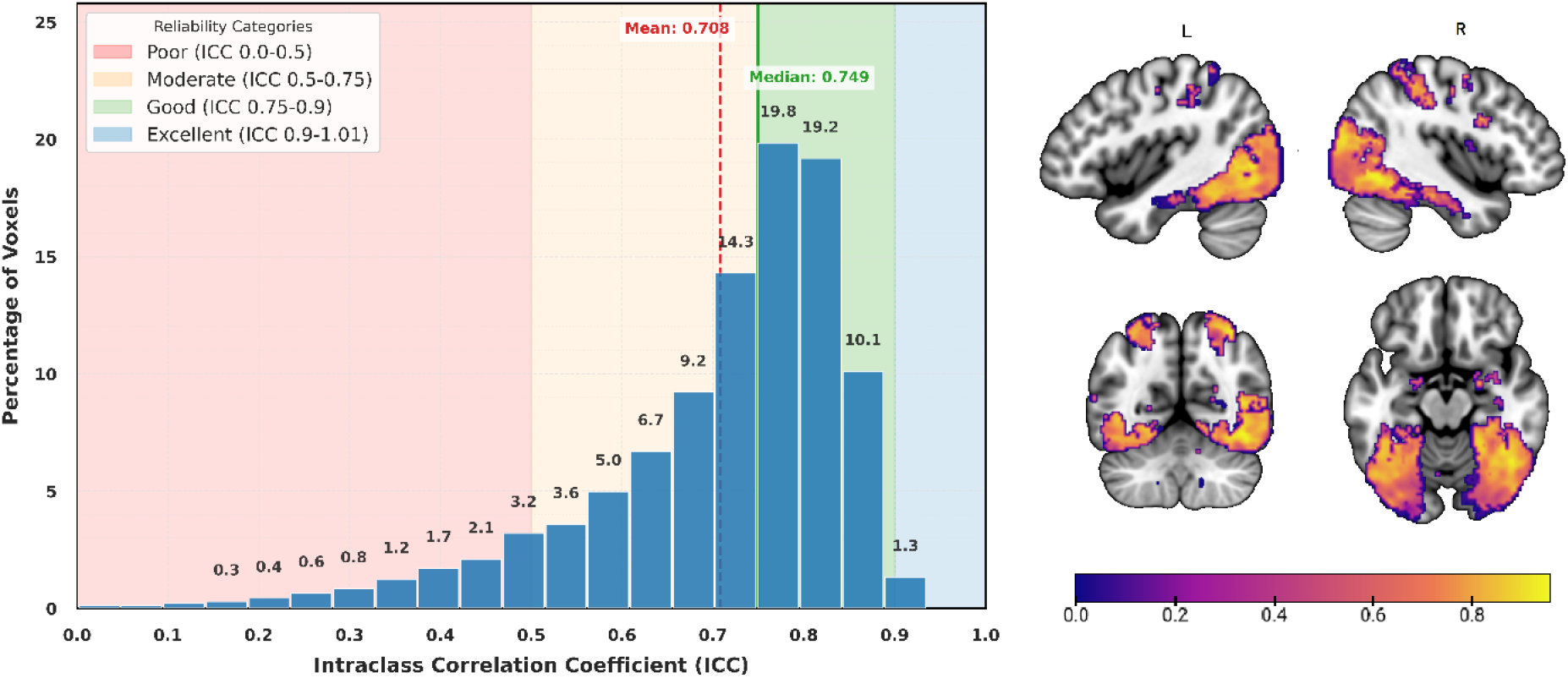
Test-retest reliability within the right Occipito-Temporal Cortex (rOTC) functional network. (Left) Histogram of voxel-wise Intraclass Correlation Coefficient (ICC) values within the rOTC network (N = 16,067 voxels). The x-axis represents ICC values binned into 20 intervals; the y-axis shows the percentage of voxels within each bin. The percentage value atop each column indicates the proportion of voxels within that specific ICC bin. Colored backgrounds denote established reliability tiers: poor (ICC < 0.50), moderate (0.50 ≤ ICC < 0.75), good (0.75 ≤ ICC < 0.90), and excellent (ICC ≥ 0.90). The red dashed vertical line indicates the mean ICC (0.70); the solid green line indicates the median ICC (0.74). (Right) Spatial distribution of ICC values within the rOTC network. Axial, coronal, and sagittal (left and right hemisphere) views are displayed on the MNI152 template. The color bar indicates ICC magnitude, matching the scale of the histogram. Voxels with the highest reliability (ICC > 0.85) are predominantly localized in the fusiform gyrus and lateral occipital cortex.

The majority of voxels within the rOTC network demonstrated good-to-excellent reliability: 72.0% of voxels showed good or excellent reliability (ICC ≥ 0.75), while 25.1% showed moderate reliability (0.50 ≤ ICC < 0.75), and only 2.9% demonstrated poor reliability (ICC < 0.50). The highest reliability values (ICC > 0.85) were predominantly observed in regions most strongly connected to the rOTC seed, including the fusiform gyrus and lateral occipital cortex.

#### 3.2.2. Reliability of the lvIFG Functional Network

A similar pattern of high reliability was observed for the lvIFG network (17,216 voxels) (**Figure 4**). The distribution of ICC values was shifted toward the high end of the reliability spectrum, with a mean of 0.74 (SD = 0.11) and a median of 0.76. The 25th and 75th percentiles were 0.68 and 0.81, respectively.

**Figure 4.**
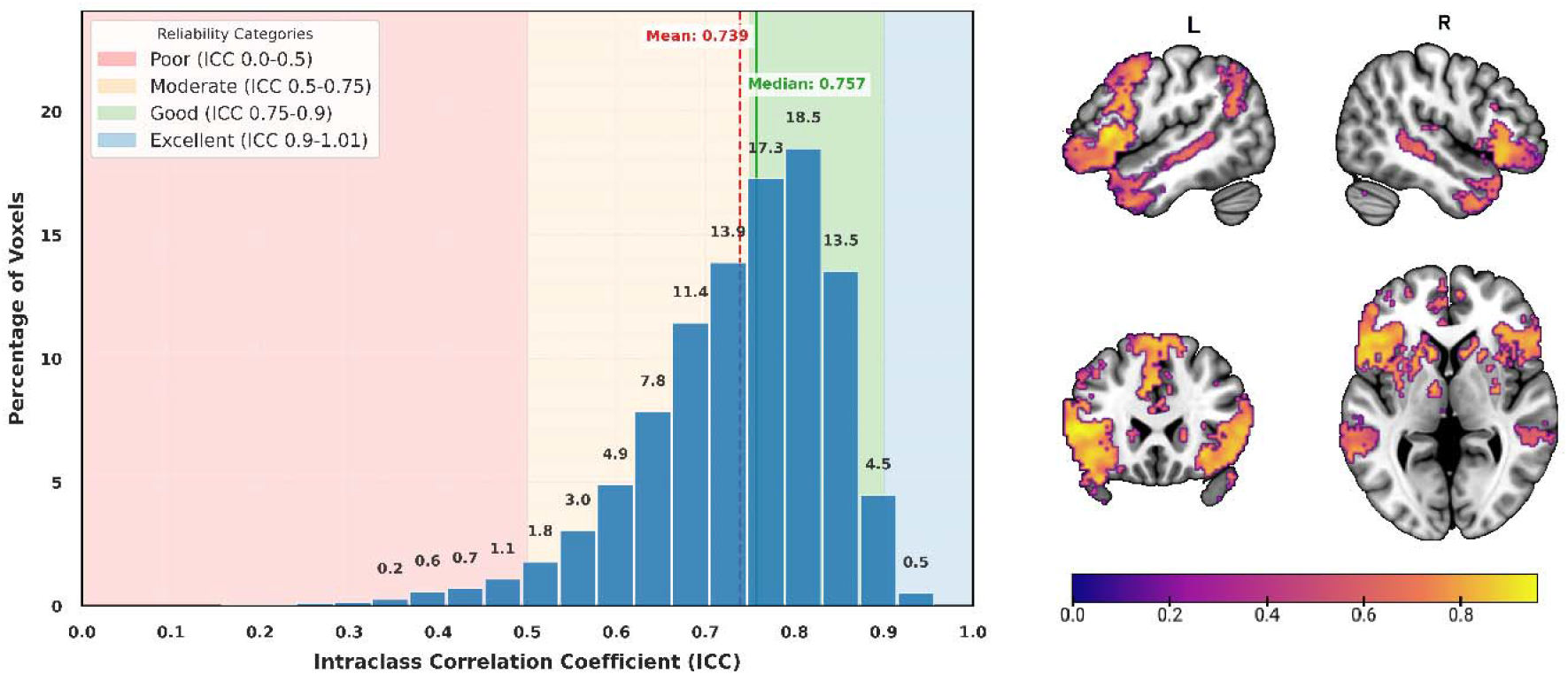
Test-retest reliability within the left Ventral Inferior Frontal Gyrus (lvIFG) functional network. (Left) Histogram of voxel-wise Intraclass Correlation Coefficient (ICC) values within the lvIFG network (N = 17,216 voxels). The x-axis represents ICC values binned into 20 intervals; the y-axis shows the percentage of voxels within each bin. The percentage value atop each column indicates the proportion of voxels within that specific ICC bin. Colored backgrounds denote established reliability tiers (Koo & Li, 2016): poor (ICC < 0.50), moderate (0.50 ≤ ICC < 0.75), good (0.75 ≤ ICC < 0.90), and excellent (ICC ≥ 0.90). The red dashed vertical line indicates the mean ICC (0.74); the solid green line indicates the median ICC (0.76). (Right) Spatial distribution of ICC values within the lvIFG network. Axial, coronal, and sagittal (left and right hemisphere) views are displayed on the MNI152 template. The color bar indicates ICC magnitude, matching the scale of the histogram. The network demonstrates a pronounced shift towards higher reliability, with the highest values concentrated in the core regions of the language network.

77.3% of voxels within the lvIFG network demonstrated good or excellent reliability (ICC ≥ 0.75). Moderate reliability was observed in 20.5% of voxels, and only 2.2% showed poor reliability. This confirms that the core functional architecture identified by the lvIFG seed is highly consistent across sessions.

### 3.3. Task-based fMRI

#### 3.3.1. Neural Correlates of Object-Location Memory

Whole-brain analysis revealed robust neural activity during the OLM learning task compared to the implicit baseline (learning > implicit baseline), engaging a widespread network including left medial temporal, fronto-parietal, and bilateral subcortical regions, consistent with the demands of OLM task (see **Supplementary Table 3A** and **Figure 5A**). The most extensive activation was a large left-lateralized cluster encompassing the parahippocampal gyrus and superior parietal lobule (SPL).

**Figure 5.**
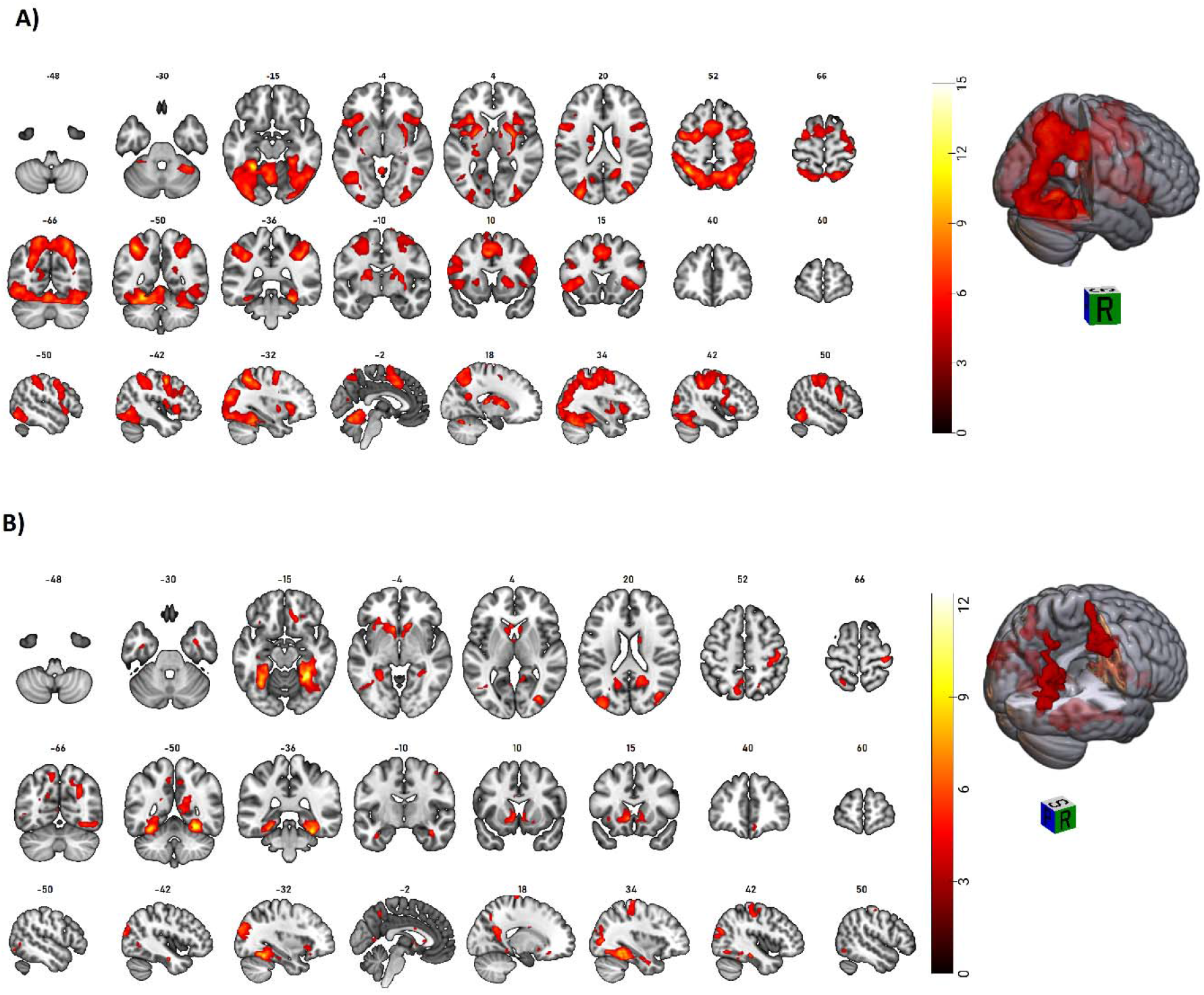
Whole-brain activation maps for the Object-Location Memory (OLM) task. (A) Learning > Implicit Baseline contrast. (B) Learning > Control contrast. All clusters are significant (pFWE < 0.05, cluster-level corrected). The color bar represents the T-score.

The direct comparison to the control condition (Learning > Control) highlighted a more specific pattern of activation, with the strongest effects observed bilaterally in the ventral visual stream, including the temporal occipital fusiform cortex and parahippocampal gyrus. Further significant activation was identified in the lateral occipital cortex, right sensorimotor cortex, bilateral precuneus, and subcortical structures (see **Supplementary Table 3B** and **Figure 5B**).

#### 3.3.2. Neural Correlates of Associative Picture-Pseudoword Learning

Whole-brain analysis revealed widespread neural activity during the APPL learning task compared to the implicit baseline (Learning > Implicit Baseline), encompassing lateral occipital, fusiform, Lingual, superior parietal, and insular regions, consistent with the demands of associative novel-word learning (see **Supplementary Table 4A** and **Figure 6A**). The most pronounced activations were found in a large bilateral cerebellar cluster, a left-dominant fronto-parietal, and temporal occipital networks

**Figure 6.**
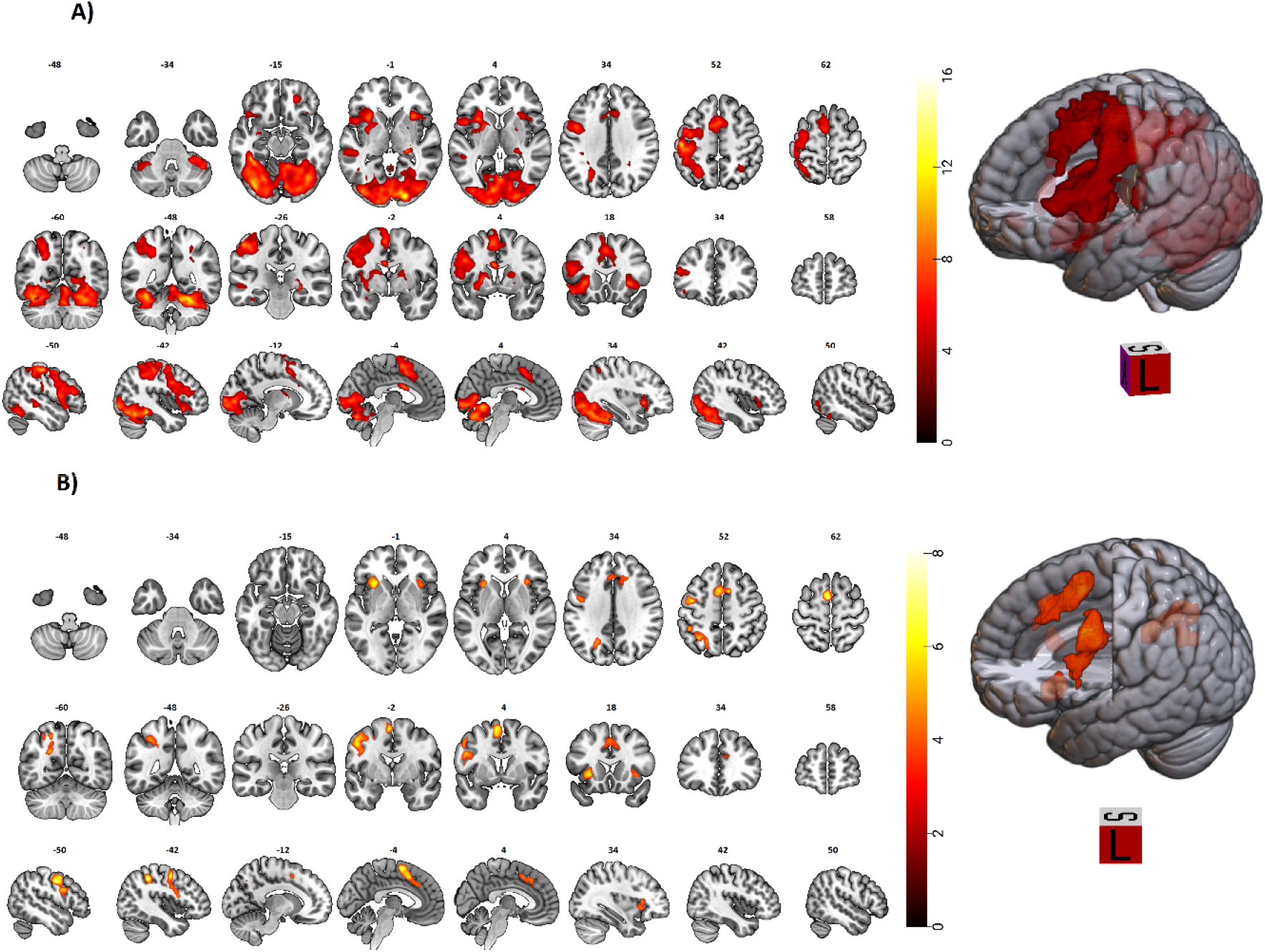
Whole-brain activation maps for the Associative Picture-Pseudoword Learning (APPL) task. (A) Learning > Implicit Baseline contrast. (B) Learning > Control contrast. All clusters are significant (pFWE < 0.05, cluster-level corrected). The color bar represents the T-score.

The direct comparison to the control condition (Learning > Control) identified a more focused set of regions specifically implicated in successful encoding. The strongest effects were found in bilateral cognitive control regions, including the SMA and paracingulate gyrus, the bilateral insular cortex, and the left precentral gyrus. (see **Supplementary Table 4B** and **Figure 6B**)

### 3.4. Overlap between Intrinsic Functional Networks and Task-Evoked Activation

#### 3.4.1. Convergence of the intrinsic rOTC Network with OLM Task

We quantified the spatial correspondence between the intrinsic rOTC network and task-evoked activity during the OLM task. A significant positive spatial correlation was observed for both the OLM > Control contrast (r = 0.289, p < 0.001) and the OLM > Baseline contrast (r = 0.365, p < 0.001). This indicates that voxels exhibiting higher task-evoked activation also demonstrated stronger intrinsic functional connectivity with the rOTC seed.

The overlap of supra-threshold voxels was substantial. For the OLM > Control contrast (see **Figure 7A**), 1,571 out of 4,198 task-activated voxels overlapped with the 8,426-voxel rOTC network (Dice = 0.249).

**Figure 7.**
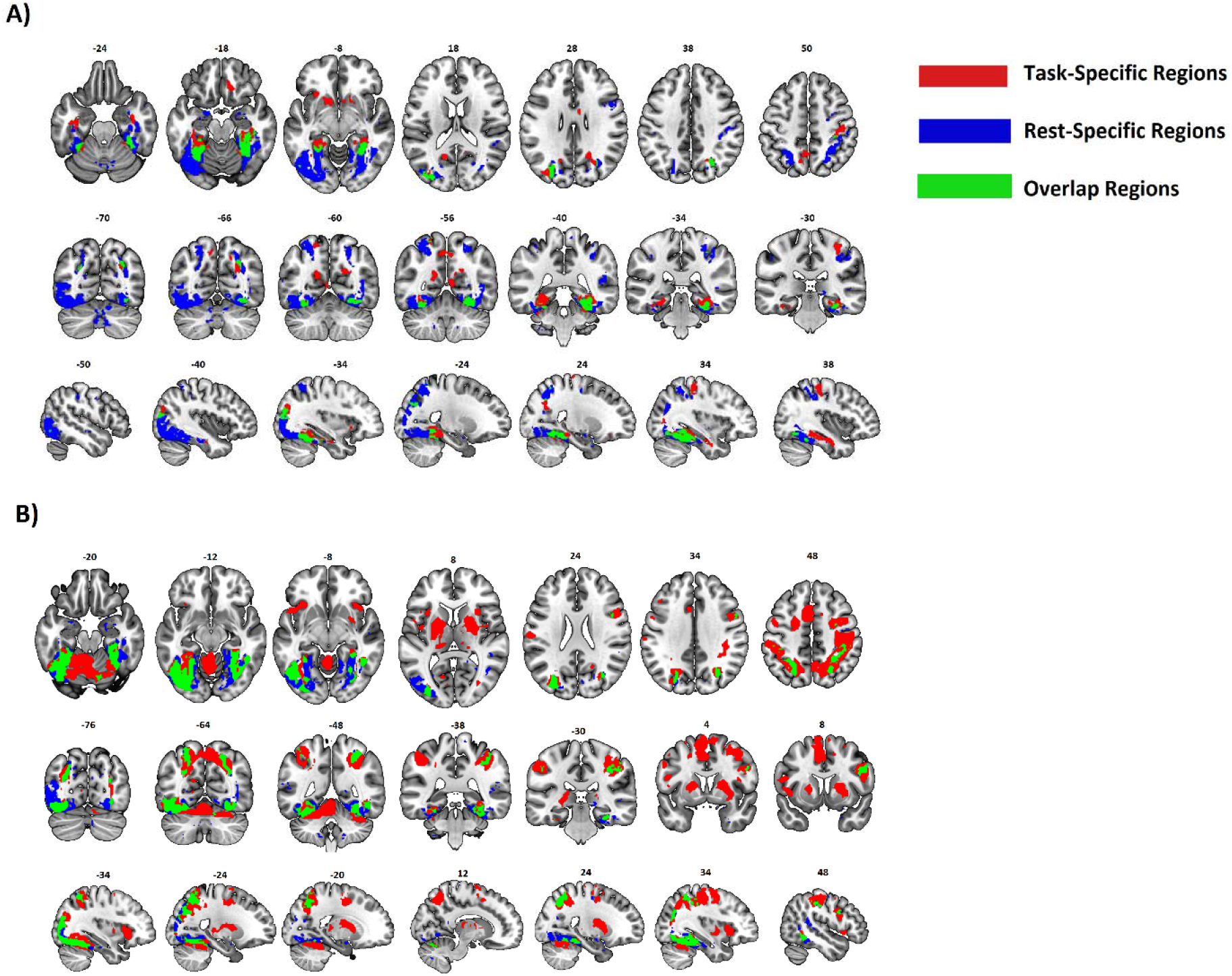
Distinct and shared neural substrates for the OLM task and rOTC network. Multi-planar views show the spatial relationship for the (A) Learning > Control and (B) Learning > Baseline contrasts. Red: task-specific activation (FWE-corrected). Blue: resting-state connectivity with the rOTC seed (FWE-corrected). Green: overlap, indicating areas both task-activated and functionally connected to the rOTC.

A notably greater overlap was found for the OLM > Baseline contrast, with 4,639 out of 18,133 task-activated voxels falling within the rOTC network (Dice = 0.349). The spatial distribution of these overlapping regions is visualized in **Figure 7B**.

#### 3.4.2. Convergence of the intrinsic lvIFG Network with APPL Task

We quantified the spatial correspondence between the intrinsic lvIFG network and task-evoked activity during the APPL task. A significant positive spatial correlation was observed for both the APPL > Control contrast (r = 0.092, p <0.001) and the APPL > Baseline contrast (r = 0.036, p < 0.001).

The overlap of supra-threshold voxels was more limited. For the APPL > Control contrast (see **Figure 8A**), 407 out of 1,382 task-activated overlapped with the 11,082-voxel lvIFG network (Dice = 0.065). For the APPL > Baseline contrast (see **Figure 8B**), 1,479 out of 16,853 task-activated voxels fell within the lvIFG network (Dice = 0.106).

**Figure 8.**
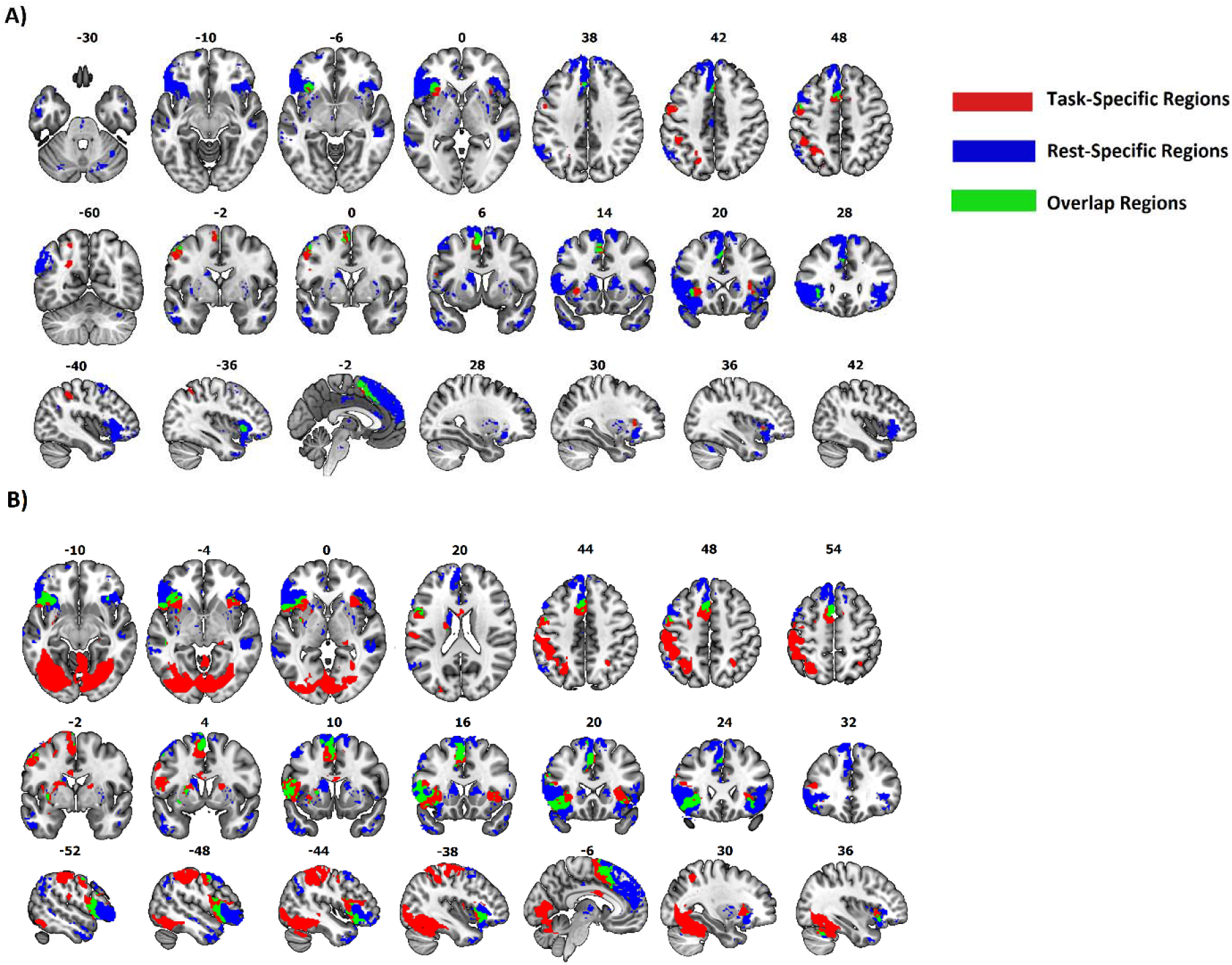
Distinct and shared neural substrates for the APPL task and lvIFG network. Multi-planar views show the spatial relationship for the (A) Learning > Control and (B) Learning > Baseline contrasts. Red: task-specific activation (FWE-corrected). Blue: resting-state connectivity with the lvIFG seed (FWE-corrected). Green: overlap, indicating areas both task-activated and functionally connected to the lvIFG.

## 4. Discussion

We describe a cross-modal fMRI-based framework to guide planning of network-specific neuromodulation and empirically guided hypothesis testing in focal tDCS-fMRI studies. We illustrated this approach by using two datasets from the MeMoSLAP consortium, that acquired both RS- and task-based fMRI at the same scanner and with identical acquisition parameters. (1) RS-data delineated the functional networks originating from the envisaged target regions for focal tDCS (rOTC, lvIFG) for two distinct experimental learning paradigms (OLM, APPL). The identified RS networks showed substantial consistency across scanning sessions, with good-to-excellent ICCs in well over 70% of voxels of either network. These findings align with long-term intrinsic connectivity networks reproducibility studies reporting ICC>0.60 for >70% of connectivity elements (Chou et al., 2012; Guo et al., 2012). This highlights the usability of seed-based RS-fMRI approaches for network-based targeting and hypothesis testing. (2) Subsequently, we determined the spatial overlap of the identified seeded-RS networks with activity elicited by the respective learning paradigms. This step identified regions in the task-active networks that could be affected by focal tDCS, either via direct application of current flow to the target region or downstream effects on functionally connected areas. Notably, only a subset of voxels identified by the seed-based RS analysis overlapped with activity elicited by either paradigm (ranging from 7.5-55%), with overall larger correspondence for the OLM task. Therefore, the degree of potential tDCS network effects varied substantially depending on the selected target region, the extent of its functional network and also task-specific activity patterns. (3) Finally, the degree of correspondence between imaging modalities was in part mediated by the chosen contrasts-of-interest in the task-based analyses, with more conservative control conditions resulting in reduced overlap between task-related activity and the seeded RS-network. This suggests that the choice of contrast has direct implications for revealing potential stimulation effects in tDCS-fMRI studies. In sum, the methodological approach exemplified in this study can be used for target network selection in focal NIBS studies and inform tDCS-fMRI data analysis and interpretation. Below, we will further discuss this novel approach and potential future extensions.

Contemporary systems neuroscience research has highlighted that higher order human brain functions rely on complex interactions between specialized brain regions, that are organized in widely distributed and partly overlapping networks (Behrens & Sporns, 2012; Breakspear, 2017). However, decisions about target selection in NIBS studies typically relied on assumptions about local processing capabilities of specific brain regions, while the origin of remote stimulation effects on brain function are currently not well understood (Meinzer et al., 2024). Notably, even some of the earliest tDCS-fMRI studies have suggested that tDCS effects may be expressed specifically within the networks of brain regions directly affected by the induced current. For example, Meinzer et al. (2012) administered conventional tDCS to left IFG during RS-fMRI and a verbal fluency task. During task-based imaging, tDCS selectively modulated activity at the stimulation site. However, in the unconstrained RS condition, tDCS increased connectivity in bilateral fronto-temporal, parietal, and premotor regions. Moreover, regions showing enhanced connectivity during RS-fMRI almost completely overlapped with two distinct networks originating from the ventral and dorsal IFG subportions, that were both directly affected by the current. These findings suggested that seed-based RS analysis may reveal areas that are potentially affected by tDCS via functional (and structural) connections and without the current reaching the connected area, thereby supporting the notion of network-based neurostimulation.

Network specific targeting is not an entirely novel approach and has recently been suggested in different contexts (Siddiqi et al., 2020; Piper et al., 2022), albeit not by using multimodal imaging. For example, a seminal study by Fox et al. (2014) employed RS-fMRI data to explore the functional connectivity between stimulation targets used in previous NIBS (cortical) and deep brain stimulation (DBS, subcortical) studies in different neuropsychiatric populations. This study confirmed that effective NIBS studies had used targets that were functionally connected to subcortical pathology foci, thereby supporting the notion of a RS network-based targeting. Likewise, we assume that neural network effects of focal tDCS are primarily expressed in brain regions that are directly connected to the target regions of NIBS, and that those can be identified by seed-based RS analysis. However, the novelty of our approach rests on the incorporation of co-acquired task-based imaging, that allowed us to determine the overlap of activity by the respective tasks with the identified seed-networks. This analysis suggested that potential tDCS effects critically depend on the interaction between the target region, its seeded network, and activity elicited by the respective experimental tasks. From a planning perspective, this approach allows to determine RS-based target networks for focal NIBS to ensure overlap with theoretically or empirically derived local nodes or networks of interest, relevant for specific tasks like OLM and APPL (e.g., Postma et al., 2008; Tagarelli et al., 2019; Abdelmotaleb et al., 2025; Kocataş et al., 2025) Moreover, our approach can also be used to determine valid active control stimulation conditions, showing limited overlap with the task-relevant networks of interest, thereby highlighting its broad applicability.

Noteworthy, our analyses revealed substantial variability of correspondence between the seeded-networks and task-based activity, ranging from 7-55%. The overlap between RS and task-based networks was most pronounced for the OLM task, which emphasizes the potential of the OTC target region to modulate activity in a larger neural network that includes core regions of the visual-limbic network underlying spatial memory formation (Abdelmotaleb et al., 2025; Postma et al., 2008) However, even in the task condition with the least degree of overlap with the corresponding seed network (i.e., the complex contrast comparing APPL with a lexical decision task), a number of task-critical regions were identified, including prefrontal, premotor and insular cortices and basal ganglia (Sliwinska et al., 2017; Tagarelli et al., 2019; Kocataş et al., 2025) These findings validate the choice of the lvIFG target region for this specific task. Based on our results, similar variability may be expected for other task- and seed-based networks and researchers are advised (a) to make informed decision about the functional relevance of potentially affected regions when choosing stimulation targets and, (b) in specific instances, limited convergence of RS- and task-activity may be trumped by the exceptional functional relevance of the few identified regions for specific processes (e.g., APPL task). Finally, both intrinsic and task-based networks are known to undergo changes across the human lifespan and even more so, in neuropsychiatric diseases (Greicius, 2008; Zhang et al., 2021). This needs to be taken into account in future studies planning to adopt a similar network-based targeting approach, ensuring that regions and networks of interests are derived from the target populations (older vs young adults; patient populations vs healthy individuals).

Our approach also allows constraining data analyses in tDCS-fMRI studies to brain regions where neural network effects of NIBS are to be expected, thereby providing an empirically informed approach for hypothesis testing. Unlike task-based imaging, RS-fMRI data is readily available from publicly available databases, including lifespan data to account for neural network reorganization due to aging (e.g., www.humanconnectome.org). This makes our framework attractive for prospective network-based targeting approaches and data analysis, with potential to rule out alternative explanations for behavioral or neural modulation like current spread to unintended target regions (Ito et al., 2017). Nonetheless, complex cognitive tasks (like OLM or APPL) typically involve several spatially or temporally distinct networks and effective processing relies on the delicate and time varying balance between them (Ren et al., 2017; Williams et al., 2022; Zhang et al., 2023). Such potential effects also need to be taken into account when planning and analyzing future tDCS-fMRI studies. Finally, our approach can be further refined by incorporating individualized current modeling (Niemann, et al., 2024) and more realistic seed regions, representing areas where (focal) tDCS induces a physiologically relevant current dose. This would allow to further constrain the networks that are directly affected by NIBS and to increase our understanding of the neural mechanisms underlying neuromodulation.

## Conclusion

This study established a principled, cross-modal fMRI framework that bridges a critical gap in neuromodulation research. By integrating reliable intrinsic connectivity maps with task-evoked activity patterns, we provide a method to prospectively identify network-level targets for focal NIBS and to generate hypotheses for data analysis in tDCS-fMRI studies. This approach paves the way for investigating the causal modulation of specific functional networks, with potential to facilitate a shift from stimulating single brain regions to targeting predefined functional pathways.

## Acknowledgments

This research was funded by the German Research Foundation (DFG) (project grants: Research Unit5429/1 [467143400]; CRC1315-B03 [327654276]; FL379/22-1; FL379/26-1; FL379/34-1; FL379/35-1; FL379/37-1; ME3161/5-1; ME3161/6-1; AN1103/5-1; Heisenberg grant to DA AN1103/6-1 [539593253]). We are grateful to our research coordinators and student assistants for their help with scheduling and testing.

## Ethics statement

The study was conducted at University Medicine Greifswald and received ethical approval from the institution’s medical ethics committee (approval number BB015/22). All procedures adhered to the Declaration of Helsinki, and written informed consent was obtained from all participants prior to enrolment.

## Data and code availability

The study was pre-registered with the Open Science Framework, and the protocol can be accessed through OSF Registries (https://osf.io/t37u2). To ensure full transparency and reproducibility, our analysis scripts are publicly available on GitHub (https://github.com/ShahAliR/NetworkOverlap-Task-RS-fMRI). Due to ethical and privacy concerns about the study’s participants, the row data are not publicly accessible. However, the corresponding author can provide the data supporting the study’s conclusions upon request.

## References

Abdelmotaleb, M., Niemann, F., Kocataş, H., Caisachana Guevara, L. M., Shahbabaie, A., Malinowski, R., Riemann, S., Fromm, A. E., Hayek, D., Antonenko, D., Meinzer, M., & Flöel, A. (2025). Identification of Reliable Target Brain Regions for Enhancing Object–Location Memory by Brain Stimulation. Brain and Behavior, 15(7), e70658. 10.1002/brb3.70658

Abraham, A., Pedregosa, F., Eickenberg, M., Gervais, P., Mueller, A., Kossaifi, J., Gramfort, A., Thirion, B., & Varoquaux, G. (2014). Machine learning for neuroimaging with scikit-learn. Frontiers in Neuroinformatics, 8. 10.3389/fninf.2014.00014

Antonenko, D., Külzow, N., Sousa, A., Prehn, K., Grittner, U., & Flöel, A. (2018). Neuronal and behavioral effects of multi-day brain stimulation and memory training. Neurobiology of Aging, 61, 245–254. 10.1016/j.neurobiolaging.2017.09.017

Behrens, T. E., & Sporns, O. (2012). Human connectomics. Current Opinion in Neurobiology, 22(1), 144– 153. 10.1016/j.conb.2011.08.005

Breakspear, M. (2017). Dynamic models of large-scale brain activity. Nature Neuroscience, 20(3), 340– 352. 10.1038/nn.4497

Chou, Y. -h., Panych, L. P., Dickey, C. C., Petrella, J. R., & Chen, N-k. (2012). Investigation of Long-Term Reproducibility of Intrinsic Connectivity Network Mapping: A Resting-State fMRI Study. American Journal of Neuroradiology, 33(5), 833–838. 10.3174/ajnr.A2894

Cirillo, G., Di Pino, G., Capone, F., Ranieri, F., Florio, L., Todisco, V., Tedeschi, G., Funke, K., & Di Lazzaro, V. (2017). Neurobiological after-effects of non-invasive brain stimulation. Brain Stimulation, 10(1), 1–18. 10.1016/j.brs.2016.11.009

De Sousa, A. V. C., Grittner, U., Rujescu, D., Külzow, N., & Flöel, A. (2020). Impact of 3-Day Combined Anodal Transcranial Direct Current Stimulation-Visuospatial Training on Object-Location Memory in Healthy Older Adults and Patients with Mild Cognitive Impairment. Journal of Alzheimer’s Disease, 75(1), 223–244. 10.3233/JAD-191234

Esmaeilpour, Z., Shereen, A. D., Ghobadi-Azbari, P., Datta, A., Woods, A. J., Ironside, M., O’Shea, J., Kirk, U., Bikson, M., & Ekhtiari, H. (2020). Methodology for tDCS integration with fMRI. Human Brain Mapping, 41(7), 1950–1967. 10.1002/hbm.24908

Esteban, O., Markiewicz, C. J., Blair, R. W., Moodie, C. A., Isik, A. I., Erramuzpe, A., Kent, J. D., Goncalves, M., DuPre, E., Snyder, M., Oya, H., Ghosh, S. S., Wright, J., Durnez, J., Poldrack, R. A., & Gorgolewski, K. J. (2019). fMRIPrep: A robust preprocessing pipeline for functional MRI. Nature Methods, 16(1), 111–116. 10.1038/s41592-018-0235-4

Fedorenko, E., Duncan, J., & Kanwisher, N. (2012). Language-Selective and Domain-General Regions Lie Side by Side within Broca’s Area. Current Biology, 22(21), 2059–2062. 10.1016/j.cub.2012.09.011

Flöel, A., Suttorp, W., Kohl, O., Kürten, J., Lohmann, H., Breitenstein, C., & Knecht, S. (2012). Non-invasive brain stimulation improves object-location learning in the elderly. Neurobiology of Aging, 33(8), 1682–1689. 10.1016/j.neurobiolaging.2011.05.007

Fox, M. D., Buckner, R. L., Liu, H., Chakravarty, M. M., Lozano, A. M., & Pascual-Leone, A. (2014). Restingstate networks link invasive and noninvasive brain stimulation across diverse psychiatric and neurological diseases. Proceedings of the National Academy of Sciences, 111(41). 10.1073/pnas.1405003111

Friston, K. J., Holmes, A. P., Worsley, K. J., Poline, J. -P., Frith, C. D., & Frackowiak, R. S. J. (1994). Statistical parametric maps in functional imaging: A general linear approach. Human Brain Mapping, 2(4), 189–210. 10.1002/hbm.460020402

Gbadeyan, O., McMahon, K., Steinhauser, M., & Meinzer, M. (2016). Stimulation of dorsolateral prefrontal cortex enhances adaptive cognitive control: A high-definition transcranial direct current stimulation study. The Journal of Neuroscience, 36(50), 12530–12536. 10.1523/JNEUROSCI.2450-16.2016

Gbadeyan, O., Steinhauser, M., McMahon, K., & Meinzer, M. (2016). Safety, Tolerability, Blinding Efficacy and Behavioural Effects of a Novel MRI-Compatible, High-Definition tDCS Set-Up. Brain Stimulation, 9(4), 545–552. 10.1016/j.brs.2016.03.018

Ghobadi-Azbari, P., Jamil, A., Yavari, F., Esmaeilpour, Z., Malmir, N., Mahdavifar-Khayati, R., Soleimani, G., Cha, Y.-H., Shereen, A. D., Nitsche, M. A., Bikson, M., & Ekhtiari, H. (2021). fMRI and transcranial electrical stimulation (tES): A systematic review of parameter space and outcomes. Progress in Neuro-Psychopharmacology and Biological Psychiatry, 107, 110149. 10.1016/j.pnpbp.2020.110149

Gillis, M. M., Garcia, S., & Hampstead, B. M. (2016). Working memory contributes to the encoding of object location associations: Support for a 3-part model of object location memory. Behavioural Brain Research, 311, 192–200. 10.1016/j.bbr.2016.05.037

Greicius, M. (2008). Resting-state functional connectivity in neuropsychiatric disorders. Current Opinion in Neurology, 21(4), 424–430. 10.1097/WCO.0b013e328306f2c5

Guo, C. C., Kurth, F., Zhou, J., Mayer, E. A., Eickhoff, S. B., Kramer, J. H., & Seeley, W. W. (2012). One-year test–retest reliability of intrinsic connectivity network fMRI in older adults. NeuroImage, 61(4), 1471–1483. 10.1016/j.neuroimage.2012.03.027

Hunold, A., Haueisen, J., Nees, F., & Moliadze, V. (2023). Review of individualized current flow modeling studies for transcranial electrical stimulation. Journal of Neuroscience Research, 101(4), 405–423. 10.1002/jnr.25154

Ito, T., Kulkarni, K. R., Schultz, D. H., Mill, R. D., Chen, R. H., Solomyak, L. I., & Cole, M. W. (2017). Cognitive task information is transferred between brain regions via resting-state network topology. Nature Communications, 8(1), 1027. 10.1038/s41467-017-01000-w

Kang, J., Lee, H., Yu, S., Lee, M., Kim, H. J., Kwon, R., Kim, S., Fond, G., Boyer, L., Rahmati, M., Koyanagi, A., Smith, L., Nehs, C. J., Kim, M. S., Sánchez, G. F. L., Dragioti, E., Kim, T., & Yon, D. K. (2024). Effects and safety of transcranial direct current stimulation on multiple health outcomes: An umbrella review of randomized clinical trials. Molecular Psychiatry, 29(12), 3789–3801. 10.1038/s41380-024-02624-3

Kocataş, H., Abdelmotaleb, M., Caisachana Guevara, L. M., Niemann, F., Shahbabaie, A., Malinowski, R., Riemann, S., Hayek, D., Antonenko, D., Rodriguez-Fornells, A., Flöel, A., & Meinzer, M. (2025). Functionally Relevant and Reliable Brain Stimulation Targets for Enhancement of Novel Word-Learning. bioRxiv preprint. 10.1101/2025.11.04.686295

Kravitz, D. J., Saleem, K. S., Baker, C. I., Ungerleider, L. G., & Mishkin, M. (2013). The ventral visual pathway: An expanded neural framework for the processing of object quality. Trends in Cognitive Sciences, 17(1), 26–49. 10.1016/j.tics.2012.10.011

Lefebvre, S., Jann, K., Schmiesing, A., Ito, K., Jog, M., Schweighofer, N., Wang, D. J. J., & Liew, S.-L. (2019). Differences in high-definition transcranial direct current stimulation over the motor hotspot versus the premotor cortex on motor network excitability. Scientific Reports, 9(1), 17605. 10.1038/s41598-019-53985-7

Lim, M., Kim, D. J., Nascimento, T. D., & DaSilva, A. F. (2024). High-definition tDCS over primary motor cortex modulates brain signal variability and functional connectivity in episodic migraine. Clinical Neurophysiology, 161, 101–111. 10.1016/j.clinph.2024.02.012

Lv, Y., Wu, S., Nitsche, M. A., Yue, T., Zschorlich, V. R., & Qi, F. (2024). A meta-analysis of the effects of transcranial direct current stimulation combined with cognitive training on working memory in healthy older adults. Frontiers in Aging Neuroscience, 16, 1454755. 10.3389/fnagi.2024.1454755

Martin, A. K., Kessler, K., Cooke, S., Huang, J., & Meinzer, M. (2020). The right temporoparietal junction Is causally associated with embodied perspective-taking. The Journal of Neuroscience, 40(15), 3089–3095. 10.1523/JNEUROSCI.2637-19.2020

Meinzer, M., Antonenko, D., Lindenberg, R., Hetzer, S., Ulm, L., Avirame, K., Flaisch, T., & Flöel, A. (2012). Electrical Brain Stimulation Improves Cognitive Performance by Modulating Functional Connectivity and Task-Specific Activation. The Journal of Neuroscience, 32(5), 1859–1866. 10.1523/JNEUROSCI.4812-11.2012

Meinzer, M., Shahbabaie, A., Antonenko, D., Blankenburg, F., Fischer, R., Hartwigsen, G., Nitsche, M. A., Li, S.-C., Thielscher, A., Timmann, D., Waltemath, D., Abdelmotaleb, M., Kocataş, H., Caisachana Guevara, L. M., Batsikadze, G., Grundei, M., Cunha, T., Hayek, D., Turker, S., … Flöel, A. (2024). Investigating the neural mechanisms of transcranial direct current stimulation effects on human cognition: Current issues and potential solutions. Frontiers in Neuroscience, 18, 1389651. 10.3389/fnins.2024.1389651

Müller, D., Habel, U., Brodkin, E. S., Clemens, B., & Weidler, C. (2023). HD-tDCS induced changes in resting-state functional connectivity: Insights from EF modeling. Brain Stimulation, 16(6), 1722– 1732. 10.1016/j.brs.2023.11.012

Narmashiri, A., & Akbari, F. (2025). The Effects of Transcranial Direct Current Stimulation (tDCS) on the Cognitive Functions: A Systematic Review and Meta-analysis. Neuropsychology Review, 35(1), 126–152. 10.1007/s11065-023-09627-x

Niemann, F., Riemann, S., Hubert, A.-K., Antonenko, D., Thielscher, A., Martin, A. K., Unger, N., Flöel, A., & Meinzer, M. (2024). Electrode positioning errors reduce current dose for focal tDCS set-ups: Evidence from individualized electric field mapping. Clinical Neurophysiology, 162, 201–209. 10.1016/j.clinph.2024.03.031

Niemann, F., Shahbabaie, A., Paßmann, S., Riemann, S., Malinowski, R., Kocataş, H., Caisachana Guevara, L. M., Abdelmotaleb, M., Antonenko, D., Blankenburg, F., Fischer, R., Hartwigsen, G., Li, S.-C., Nitsche, M. A., Thielscher, A., Timmann, D., Fromm, A., Hayek, D., Hubert, A.-K., … Meinzer, M. (2024). Neuronavigated Focalized Transcranial Direct Current Stimulation Administered During Functional Magnetic Resonance Imaging. Journal of Visualized Experiments, 213, 67155. 10.3791/67155

Oldfield, R. C. (1971). The assessment and analysis of handedness: The Edinburgh inventory. Neuropsychologia, 9(1), 97–113. 10.1016/0028-3932(71)90067-4

Opitz, A., Yeagle, E., Thielscher, A., Schroeder, C., Mehta, A. D., & Milham, M. P. (2018). On the importance of precise electrode placement for targeted transcranial electric stimulation. NeuroImage, 181, 560–567. 10.1016/j.neuroimage.2018.07.027

Ostrowski, J., Svaldi, J., & Schroeder, P. A. (2022). More focal, less heterogeneous? Multi-level meta-analysis of cathodal high-definition transcranial direct current stimulation effects on language and cognition. Journal of Neural Transmission, 129(7), 861–878. 10.1007/s00702-022-02507-3

Perceval, G., Martin, A. K., Copland, D. A., Laine, M., & Meinzer, M. (2020). Multisession transcranial direct current stimulation facilitates verbal learning and memory consolidation in young and older adults. Brain and Language, 205, 104788. 10.1016/j.bandl.2020.104788

Piper, R. J., Richardson, R. M., Worrell, G., Carmichael, D. W., Baldeweg, T., Litt, B., Denison, T., & Tisdall, M. M. (2022). Towards network-guided neuromodulation for epilepsy. Brain, 145(10), 3347– 3362. 10.1093/brain/awac234

Postma, A., Kessels, R., & Vanasselen, M. (2008). How the brain remembers and forgets where things are: The neurocognition of object–location memory. Neuroscience & Biobehavioral Reviews, 32(8), 1339–1345. 10.1016/j.neubiorev.2008.05.001

Prehn, K., Stengl, H., Grittner, U., Kosiolek, R., Ölschläger, A., Weidemann, A., & Flöel, A. (2017). Effects of Anodal Transcranial Direct Current Stimulation and Serotonergic Enhancement on Memory Performance in Young and Older Adults. Neuropsychopharmacology, 42(2), 551–561. 10.1038/npp.2016.170

Ren, S., Li, J., Taya, F., deSouza, J., Thakor, N. V., & Bezerianos, A. (2017). Dynamic Functional Segregation and Integration in Human Brain Network During Complex Tasks. IEEE Transactions on Neural Systems and Rehabilitation Engineering, 25(6), 547–556. 10.1109/TNSRE.2016.2597961

Riemann, S., Van Lück, J., Rodríguez-Fornells, A., Flöel, A., & Meinzer, M. (2024). The role of frontal cortex in novel-word learning and consolidation: Evidence from focal transcranial direct current stimulation. Cortex, 177, 15–27. 10.1016/j.cortex.2024.05.004

Rolls, E. T., Zhang, R., Deco, G., Vatansever, D., & Feng, J. (2024). Selective Brain Activations and Connectivities Related to the Storage and Recall of Human Object-Location, Reward-Location, and Word-Pair Episodic Memories. Human Brain Mapping, 45(15), e70056. 10.1002/hbm.70056

Satterthwaite, T. D., Elliott, M. A., Gerraty, R. T., Ruparel, K., Loughead, J., Calkins, M. E., Eickhoff, S. B., Hakonarson, H., Gur, R. C., Gur, R. E., & Wolf, D. H. (2013). An improved framework for confound regression and filtering for control of motion artifact in the preprocessing of resting-state functional connectivity data. NeuroImage, 64, 240–256. 10.1016/j.neuroimage.2012.08.052

Saturnino, G. B., Siebner, H. R., Thielscher, A., & Madsen, K. H. (2019). Accessibility of cortical regions to focal TES: Dependence on spatial position, safety, and practical constraints. NeuroImage, 203, 116183. 10.1016/j.neuroimage.2019.116183

Setsompop, K., Gagoski, B. A., Polimeni, J. R., Witzel, T., Wedeen, V. J., & Wald, L. L. (2012). Blipped-controlled aliasing in parallel imaging for simultaneous multislice echo planar imaging with reduced g -factor penalty. Magnetic Resonance in Medicine, 67(5), 1210–1224. 10.1002/mrm.23097

Siddiqi, S. H., Taylor, S. F., Cooke, D., Pascual-Leone, A., George, M. S., & Fox, M. D. (2020). Distinct Symptom-Specific Treatment Targets for Circuit-Based Neuromodulation. American Journal of Psychiatry, 177(5), 435–446. 10.1176/appi.ajp.2019.19090915

Sliwinska, M. W., Violante, I. R., Wise, R. J. S., Leech, R., Devlin, J. T., Geranmayeh, F., & Hampshire, A. (2017). Stimulating Multiple-Demand Cortex Enhances Vocabulary Learning. The Journal of Neuroscience, 37(32), 7606–7618. 10.1523/JNEUROSCI.3857-16.2017

Tagarelli, K. M., Shattuck, K. F., Turkeltaub, P. E., & Ullman, M. T. (2019). Language learning in the adult brain: A neuroanatomical meta-analysis of lexical and grammatical learning. NeuroImage, 193, 178–200. 10.1016/j.neuroimage.2019.02.061

Vallat, R. (2018). Pingouin: Statistics in Python. Journal of Open Source Software, 3(31), 1026. 10.21105/joss.01026

Wang, H.-T., Meisler, S. L., Sharmarke, H., Clarke, N., Gensollen, N., Markiewicz, C. J., Paugam, F., Thirion, B., & Bellec, P. (2023). Continuous Evaluation of Denoising Strategies in Resting-State fMRI Connectivity Using fMRIPrep and Nilearn. bioRxiv. 10.1101/2023.04.18.537240

Williams, K. A., Numssen, O., & Hartwigsen, G. (2022). Task-specific network interactions across key cognitive domains. Cerebral Cortex, 32(22), 5050–5071. 10.1093/cercor/bhab531

Zhang, H., Gertel, V. H., Cosgrove, A. L., & Diaz, M. T. (2021). Age-related differences in resting-state and task-based network characteristics and cognition: A lifespan sample. Neurobiology of Aging, 101, 262–272. 10.1016/j.neurobiolaging.2020.10.025

Zhang, H., Meng, C., Di, X., Wu, X., & Biswal, B. (2023). Static and dynamic functional connectome reveals reconfiguration profiles of whole-brain network across cognitive states. Network Neuroscience, 7(3), 1034–1050. 10.1162/netn_a_00314

