## Supplementary Materials for "Multimodal Imaging-Based Targeting Approach for Network-Level Brain Stimulation"

| Anatomical Label (Harvard–Oxford cortical–subcortical structural atlas) | Hem. | Cluster Size (voxels) | Cluster p-FWE^*^ | Peak T- values | Peak coordinates (MNI) | | |
| --- | --- | --- | --- | --- | --- | --- | --- |
|  |  |  |  |  | **x** | **y** | **z** |
| Occipital Fusiform Gyrus (Lateral visual stream) | R | 8973 | 0 | 17.70 | 46 | -62 | -10 |
| Occipital Pole (Medial early visual) | L | 6682 | 0 | 11.19 | -42 | -88 | 6 |
| Cerebellum (Crus I) | R | 394 | 0 | 8.56 | 6 | -72 | -36 |
| Lateral Occipital Cortex (Superior division) | R | 174 | 0 | 7.10 | 52 | -44 | 12 |
| Precentral Gyrus (Dorsal premotor) | R | 170 | 0 | 5.05 | 52 | 8 | 30 |
| Supramarginal Gyrus (Posterior division - PF) | L | 153 | 0 | 5.47 | -38 | -36 | 38 |
| Superior Temporal Gyrus (Posterior auditory) | L | 82 | 0 | 5.04 | -64 | -50 | 16 |
| Temporal Fusiform Cortex (Posterior face area) | L | 65 | 0 | 6.02 | -40 | -26 | -22 |
| Brainstem (Midbrain) | - | 46 | 0 | 6.64 | 16 | -28 | -4 |
| Parahippocampal Gyrus (Anterior collateral sulcus) | R | 43 | 0 | 5.17 | 30 | -2 | -34 |
| Cerebellum (VI lobe) | L | 42 | 0.000001 | 6.10 | -16 | -52 | -48 |
| Postcentral Gyrus (Hand area - 3b) | R | 41 | 0.000001 | 5.39 | 26 | -10 | 52 |
| Amygdala (Laterobasal group) | R | 39 | 0.000002 | 6.06 | 22 | -4 | -18 |
| Amygdala (Centromedial group) | L | 35 | 0.000007 | 5.78 | -20 | -6 | -18 |
| Temporal Pole (Medial) | R |  |  | 6.41 | 34 | 0 | -16 |
| Precentral Gyrus (Ventral premotor) | R | 32 | 0.000019 | 4.55 | 40 | -12 | 46 |
| Superior Temporal Gyrus (Anterior auditory) | R |  |  | 5.79 | 56 | -16 | -4 |
| Cerebellum (VIIb) | R | 30 | 0.000039 | 4.92 | 16 | -50 | -48 |
| Postcentral Gyrus (Face area - 3b) | L | 29 | 0.000056 | 4.58 | -50 | -2 | 40 |
| Precentral Gyrus (Frontal operculum) | R | 28 | 0.00008 | 4.32 | 44 | -4 | 50 |
| Occipital Pole (Peripheral V1) | L | 27 | 0.000115 | 5.39 | -6 | -90 | 24 |
| Temporal Fusiform Cortex (Anterior word area) | L | 24 | 0.000357 | 5.98 | -50 | -10 | -10 |
| Cerebellum (VIIIa) | L | 21 | 0.00116 | 4.29 | -26 | -38 | -50 |
| Parahippocampal Gyrus (Posterior collateral sulcus) | L | 19 | 0.002625 | 5.27 | -32 | -8 | -34 |
| Temporal Pole (Lateral) | R | 18 | 0.00399 | 5.31 | 56 | 10 | -22 |
| Supramarginal Gyrus (Anterior division - PFt) | L | 16 | 0.009428 | 4.22 | -36 | -40 | 58 |
| Superior Temporal Gyrus (Middle auditory) |  |  |  | 4.79 | -56 | -28 | 2 |
| Postcentral Gyrus (Arm area - 1) | R | 13 | 0.03642 | 3.91 | 16 | -26 | 70 |
| Middle Temporal Gyrus (Temporooccipital) | L |  |  | 4.93 | -60 | 4 | -20 |
| Cerebellum (IX) | L |  |  | 4.71 | -30 | -66 | -48 |

**Supplementary Table 1**: Whole-brain resting-state functional connectivity of the right Occipito-Temporal Cortex (rOTC) seed region

*p-FWE values of 0 indicate p < 0.000001

Note: Cluster-based inference used Random Field Theory (RFT) with a voxel-wise threshold of p < 0.001 and cluster-forming threshold of p-FWE < 0.05. Reported clusters survive family-wise error correction (cluster-size FWE) with a minimum extent of 13 voxels.

| Anatomical Label (Harvard–Oxford cortical–subcortical structural atlas) | Hem. | Cluster Size (voxels) | Cluster p-FWE^*^ | Peak T- values | Peak Coordinates (MNI) | | |
| --- | --- | --- | --- | --- | --- | --- | --- |
|  |  |  |  |  | **x** | **y** | **z** |
| Superior Temporal Gyrus, Anterior Division | L | 13145 | 0 | 20.32 | -48 | 20 | 2 |
| Temporal Pole | R | 2159 | 0 | 9.26 | 52 | 26 | 0 |
| Angular Gyrus (PGp) | L | 1038 | 0 | 7.57 | -62 | -52 | 30 |
| Inferior Temporal Gyrus, Temporooccipital | R | 528 | 0 | 6.73 | 38 | 16 | -42 |
| Middle Temporal Gyrus | R | 456 | 0 | 8.29 | 62 | -36 | 0 |
| Occipital Fusiform Gyrus | R | 246 | 0 | 6.62 | 30 | -72 | -28 |
| Thalamus (Pulvinar) | L | 160 | 0 | 7.03 | -6 | -14 | 0 |
| Cingulate Gyrus, Mid-Posterior | L | 124 | 0 | 8.33 | -2 | -18 | 36 |
| Cerebellum (Crus I) | R | 99 | 0 | 6.25 | 36 | -54 | -32 |
| Anterior Cingulate Cortex | L | 76 | 0 | 7.28 | -2 | 26 | 16 |
| Supramarginal Gyrus, Posterior Division | R | 59 | 0 | 5.19 | 64 | -46 | 30 |
| Parahippocampal Gyrus | L | 38 | 0.000002 | 6.38 | -22 | 2 | -38 |
| Cerebellum (VIIIa) | L | 35 | 0.000005 | 5.92 | -28 | -74 | -30 |
| Supramarginal Gyrus, Anterior Division | L | 33 | 0.000010 | 5.13 | -32 | -32 | 18 |
| Brainstem (Midbrain) | – | 19 | 0.002083 | 6.50 | 0 | -18 | -30 |
| Hippocampus | L | 14 | 0.019202 | 7.85 | -26 | -26 | -16 |
| Precuneus Cortex | L | 16 | 0.007701 | 5.19 | -8 | -54 | 30 |
| Frontal Pole | R |  |  | 5.19 | 38 | 50 | -12 |
| Orbital Frontal Cortex | L | 13 | 0.030731 | 4.41 | -2 | 46 | -20 |
| Precentral Gyrus (Dorsal) | R | 12 | 0.049600 | 4.62 | 44 | 12 | 46 |
| Precentral Gyrus (Ventral) | R |  |  | 4.62 | 42 | 20 | 40 |
| Postcentral Gyrus | R |  |  | 4.62 | 46 | -26 | 10 |
| Caudate Nucleus | R |  |  | 4.65 | 6 | -8 | 6 |
| Thalamus (Mediodorsal) | R |  |  | 6.98 | 8 | -14 | -2 |

**Supplementary Table 2**: Whole-brain resting-state functional connectivity of the left ventral Inferior Frontal Gyrus (lvIFG) seed region.

* p-FWE values of 0 indicate p < 0.000001

Note: Cluster-based inference used Random Field Theory (RFT) with a voxel-wise threshold of p < 0.001 and cluster-forming threshold of p-FWE < 0.05. Reported clusters survive family-wise error correction (cluster-size FWE) with a minimum extent of 12 voxels.

| Anatomical Label (Harvard–Oxford cortical–subcortical structural atlas) | Hem. | Cluster Size (voxels) | Cluster p-FWE^*^ | Peak T- values | Peak coordinates (MNI) | | |
| --- | --- | --- | --- | --- | --- | --- | --- |
|  |  |  |  |  | **x** | **y** | **z** |
| Parahippocampal Gyrus | L | 20404 | 0.000 | 15.35 | -26 | -46 | -18 |
| Superior Parietal Lobule | L |  |  | 11.85 | -34 | -50 | 50 |
| Cerebellum (VI) | R |  |  | 10.91 | 28 | -40 | -18 |
| Precentral Gyrus | L | 6380 | 0.000 | 10.57 | -42 | -4 | 48 |
| Thalamus | L |  |  | 10.55 | -16 | -26 | 12 |
| Supplementary Motor Area | L |  |  | 9.98 | -6 | 8 | 42 |
| Caudate | R | 2592 | 0.000 | 10.40 | 22 | -2 | 6 |
| Thalamus | R |  |  | 9.11 | 18 | -24 | 10 |
| Caudate / Putamen | R |  |  | 8.99 | 26 | -6 | 14 |
| Precuneous Cortex | R | 217 | 0.001 | 5.48 | -8 | -70 | 14 |
| Precuneous Cortex | R |  |  | 5.29 | -16 | -64 | 20 |
| Occipital Pole | L |  |  | 4.47 | -8 | -80 | 4 |
| Precuneous Cortex | R | 205 | 0.002 | 6.45 | 18 | -56 | 22 |

**Supplementary Table 3A**: Brain regions showing significant activation for the Object-Location Memory (OLM) Learning > Implicit baseline contrast.

*p-FWE values of 0 indicate p < 0.000001

| Anatomical Label (Harvard–Oxford cortical–subcortical structural atlas) | Hem. | Cluster Size (voxels) | Cluster p-FWE^*^ | Peak T- values | Peak coordinates (MNI) | | |
| --- | --- | --- | --- | --- | --- | --- | --- |
|  |  |  |  |  | **x** | **y** | **z** |
| Temporal Occipital Fusiform Cortex | R | 1533 | 0 | 12.96 | 30 | -46 | -14 |
| Temporal Fusiform Cortex, posterior division | R |  |  | 9.60 | 30 | -36 | -22 |
| Temporal Fusiform Cortex | R |  |  | 8.77 | 40 | -26 | -18 |
| Temporal Occipital Fusiform Cortex | L | 1207 | 0 | 9.17 | -32 | -50 | -18 |
| Temporal Occipital Fusiform Cortex | L |  |  | 8.87 | -32 | -58 | -16 |
| Parahippocampal Gyrus, posterior division | L |  |  | 8.60 | -26 | -36 | -18 |
| Lateral Occipital Cortex, superior division | L | 734 | 0 | 7.79 | -36 | -80 | 26 |
| Lateral Occipital Cortex, superior division | L |  |  | 6.88 | -40 | -86 | 18 |
| Lateral Occipital Cortex, Superior | L |  |  | 5.70 | -30 | -82 | 32 |
| Lateral Occipital Cortex, Superior | R | 1272 | 0 | 7.03 | 32 | -76 | 6 |
| Lateral Occipital Cortex, Superior | R |  |  | 6.78 | 40 | -72 | 20 |
| Lateral Occipital Cortex, Superior | R |  |  | 6.49 | 26 | -64 | 32 |
| Precentral Gyrus | R | 614 | 0 | 6.51 | 16 | -24 | 76 |
| Postcentral Gyrus | R |  |  | 6.25 | 34 | -26 | 68 |
| Postcentral Gyrus | R |  |  | 5.65 | 44 | -22 | 50 |
| Accumbens | L | 526 | 0 | 7.48 | -8 | 14 | -4 |
| Caudate | L |  |  | 6.38 | -4 | 20 | 4 |
| Thalamus | L |  |  | 6.02 | 0 | -2 | 4 |
| Precuneous Cortex | R | 227 | 0.001 | 4.98 | -14 | -62 | 20 |
| Lingual Gyrus | R |  |  | 4.65 | -4 | -62 | 8 |
| Precuneous Cortex | L | 460 | 0 | 5.89 | -4 | -56 | 50 |
| Precuneous Cortex | R |  |  | 5.27 | 8 | -52 | 44 |
| Precuneous Cortex | R |  |  | 4.96 | -10 | -68 | 52 |
| Caudate | L | 185 | 0.004 | 6.53 | 12 | 20 | -4 |
| Accumbens | L |  |  | 5.26 | 6 | 10 | -6 |
| Putamen | R |  |  | 4.60 | 20 | 10 | -8 |
| Cingulate Gyrus, anterior division (posterior part) | R | 139 | 0.017 | 6.32 | 8 | 0 | 28 |
| Caudate | R |  |  | 4.97 | 14 | -2 | 22 |
| Frontal Medial Cortex | R | 157 | 0.009 | 6.34 | 12 | 32 | -18 |
| Frontal Orbital Cortex | R |  |  | 4.37 | 26 | 32 | -14 |

**Supplementary Table 3B**: Brain regions showing significant activation for the Object-Location Memory (OLM) Learning > Control contrast. Significant clusters surviving family-wise error (FWE) correction at the cluster level (p < 0.05).

*p-FWE values of 0 indicate p < 0.000001

| Anatomical Label (Harvard–Oxford cortical–subcortical structural atlas) | Hem. | Cluster Size (voxels) | Cluster p-FWE* | Peak T-value | Peak coordinates (MNI) | | |
| --- | --- | --- | --- | --- | --- | --- | --- |
|  |  |  |  |  | X | Y | Z |
| Cerebellum VI | R | 16594 | 0 | 15.72 | 24 | -52 | -20 |
| Lingual gyrus | R |  |  | 15.44 | 14 | -86 | -6 |
| Lateral occipital cortex | R |  |  | 15.10 | 22 | -92 | 0 |
| Temporal Occipital Fusiform Cortex | L |  |  | 11.32 | -40 | -58 | -19 |
| Temporal Fusiform Cortex posterior division | L |  |  | 10.96 | -39 | -40 | -19 |
| Thalamus | R | 133 | 0.005 | 11.97 | 24 | -28 | -4 |
| Pallidum | R |  |  | 7.19 | 28 | -20 | -6 |
| Postcentral gyrus | L | 10562 | 0 | 11.22 | -52 | -20 | 52 |
| Supplementary Motor Cortex | L |  |  | 9.58 | -6 | 0 | 58 |
| Putamen | L |  |  | 9.40 | -20 | 4 | 6 |
| Cingulate gyrus anterior division | L | 200 | 0.001 | 9.89 | -6 | 2 | 28 |
| Cingulate gyrus anterior division | L |  |  | 4.45 | -6 | -10 | 28 |
| Insular cortex | R | 392 | 0 | 9.34 | 32 | 18 | 0 |
| Frontal Orbital Cortex | R |  |  | 4.68 | 36 | 24 | -12 |
| Frontal Operculum Cortex | R |  |  | 4.57 | 42 | 14 | 6 |
| Frontal Orbital Cortex | R | 110 | 0.010 | 7.39 | 20 | 42 | -14 |
| Putamen | R | 181 | 0.001 | 5.65 | 18 | 0 | 12 |
| Thalamus | R |  |  | 3.74 | 20 | -12 | 6 |
| Thalamus | R |  |  | 3.45 | 16 | -16 | 12 |
| Superior parietal lobule | R | 169 | 0.002 | 4.21 | 28 | -52 | 50 |
| Lateral Occipital Cortex, superior division | R |  |  | 3.60 | 34 | -54 | 56 |

**Supplementary Table 4A**: Brain regions showing significant activation for the Associative Picture-Pseudoword Learning (APPL) Learning > Implicit baseline contrast.

* p-FWE values of 0 indicate p < 0.0001

| Anatomical Label (Harvard–Oxford cortical–subcortical structural atlas) | Hem. | Cluster Size (voxels) | Cluster p-FWE* | Peak T- values | Peak Coordinates (MNI) | | |
| --- | --- | --- | --- | --- | --- | --- | --- |
|  |  |  |  |  | **x** | **y** | **z** |
| Supramarginal Gyrus, posterior division | L | 793 | 0 | 7.43 | -44 | -42 | 48 |
| Lateral Occipital Cortex, superior division | L |  |  | 7.17 | -24 | -64 | 32 |
| Superior Parietal Lobule | L |  |  | 6.98 | -32 | -54 | 48 |
| Precentral Gyrus | L | 852 | 0 | 8.07 | -50 | -8 | 42 |
| Precentral Gyrus | L |  |  | 6.96 | -40 | -4 | 48 |
| Middle Frontal Gyrus, posterior division | L |  |  | 5.99 | -50 | 2 | 48 |
| Supplementary Motor Cortex | L | 936 | 0 | 8.04 | -4 | 0 | 62 |
| Paracingulate gyrus | R |  |  | 6.03 | 10 | 8 | 52 |
| Paracingulate gyrus | L |  |  | 4.64 | -2 | 14 | 48 |
| Insular Cortex | L | 273 | 0.001 | 8.44 | -30 | 22 | -2 |
| Insular Cortex | R | 143 | 0.017 | 8.25 | 32 | 22 | 10 |
| Insular Cortex | R |  |  | 5.99 | 36 | 18 | -2 |

**Supplementary Table 4B**: Brain regions showing significant activation for the Associative Picture-Pseudoword Learning (APPL) Learning > Control contrast. Significant clusters surviving family-wise error (FWE) correction at the cluster level (p < 0.05).

* p-FWE values of 0 indicate p < 0.0001
